# Introducing Mouffet, a unified framework to make model creation easier and more reproducible

**DOI:** 10.1101/2022.07.06.498965

**Authors:** Sylvain Christin, Nicolas Lecomte

## Abstract

1. Biological and ecological models are being increasingly used to explain the natural world. Model creation is an iterative process requiring two steps: training and evaluating the models. However, this process can become complex when multiple models are trained and evaluated at the same time. Besides, development steps can be lost, reducing the reproducibility of model creation.
2. We introduce Mouffet, an open-source Python framework that aims to make model creation easier, more robust, and more reproducible. It provides a set of configuration files and high-level Python interfaces that help managing data, training, and evaluating models. To improve reproducibility, every step of the model creation process, including the options used, are saved.
3. Mouffet introduces the notion of scenarios that allow users to define multiple training or evaluation tasks in a single configuration file. This not only facilitates model creation but enables users to define experimental plans to study the effect of selected parameters on training or evaluation.
4. While initially developed for deep learning models, Mouffet is independent of the implementation of the models. Therefore, it could be successfully used to compare different modelling approaches. Besides, its ease of use makes it a choice tool for ecologists, even when not familiar with complex model creation.

## 1. Introduction

Biological models have become an essential tool to better understand our world (Jackson et al., 2000; Shoemaker et al., 2021). They are now becoming more and more numerous and sophisticated, even more so with the recent progress made in machine learning, and especially deep learning (Borowiec et al., 2022; Christin et al., 2019; Guo et al., 2020). Such fast evolution, however, introduces challenges that make the creation and comparison of new models difficult, hampering their reproducibility and research potential.

The development of a model is an iterative process, with many candidates often created and improved upon during development (Jackson et al., 2000). Two main tasks are required to develop a model, the first being to create the model. In the machine learning world, where a model adjusts internal parameters from data, this step is called training (Webb, 2002). The second task of model development is to verify that the model does what it was created for (Christin et al., 2021; Quinn et al., 2021). Doing so, several candidates are often compared to select the best one (Webb, 2002). To choose the right model, several evaluation methods can be used. Indeed, a single model might perform well on one aspect of a problem but poorly on another. For instance, let’s say we try to create a bird song detector from 30-second audio samples. One model could prove very efficient in telling us whether a bird song is present in a sample, but less so to tell when this song occurs. This could lead to situations where we obtain multiple models that need to be evaluated with different methods.

This introduces several challenges. First, this can quickly become messy as many results need to be compiled and tracked. Then, even when a model is freely made available after publication, often only the code for the “winning” model is presented, and steps leading to its development can be lost, reducing the reproducibility of the research. At a time when science becomes more and more open access and reproducible (Baker, 2016; Sullivan et al., 2019), we believe that model creation should follow a well-defined experimental plan that can easily be run, saved, compared, and repeated.

Many tools exist to assist in model creation. They often offer ways to manage data, training, and evaluation, e.g Tensorflow (Hope et al., 2017) or Pytorch (Paszke et al., 2019) for deep learning. However, these tools are usually targeted for computer scientists. Besides, to our knowledge, none offer a rigorous, structured, and easy way to train and evaluate multiple models, while saving all the information required to reproduce these steps.

Here we introduce Mouffet, an open-source Python framework aiming at making model creation easier, more robust, and more repeatable. To do so, Mouffet provides a set of high-level interfaces and tools, easily customizable through configuration files, to facilitate and keep track of each aspect of model creation. As its design is independent of the underlying implementations of the models, Mouffet can be used for virtually any kind of model. By introducing the notion of scenarios, that allow the training or evaluation of several models within a single configuration file, ecologists can now easily define experimental plans for model creation that can easily be shared and repeated.

## 2. Mouffet

Mouffet was created to allow users to define reproducible experimental protocols for any model creation task, i.e., training or evaluation. It is based on the following principles:

- Options are easily accessible and customizable
- Results are saved in a comprehensive way
- Code is defined in a structured yet flexible way
- Multiple tasks can be performed with a single command call

To achieve its goals, Mouffet relies heavily on user-friendly configuration files. This allows all options needed during model creation to be free of the code logic while remaining quickly accessible. They are based on the YAML format, a format that is both easy to read and easy to use. Mouffet provides tools to easily access these files and the options they contain, and users can add any option they deem useful. A full description and explanation of how Mouffet works can be found at (https://mouffet.readthedocs.io/en/latest/index.html).

### 2.1 Configuration files

By default, three types of configuration files are used: one for data management, one to train the models, and one to evaluate the models (Fig. 1). Here is a brief description of each one.

**Figure 1:**
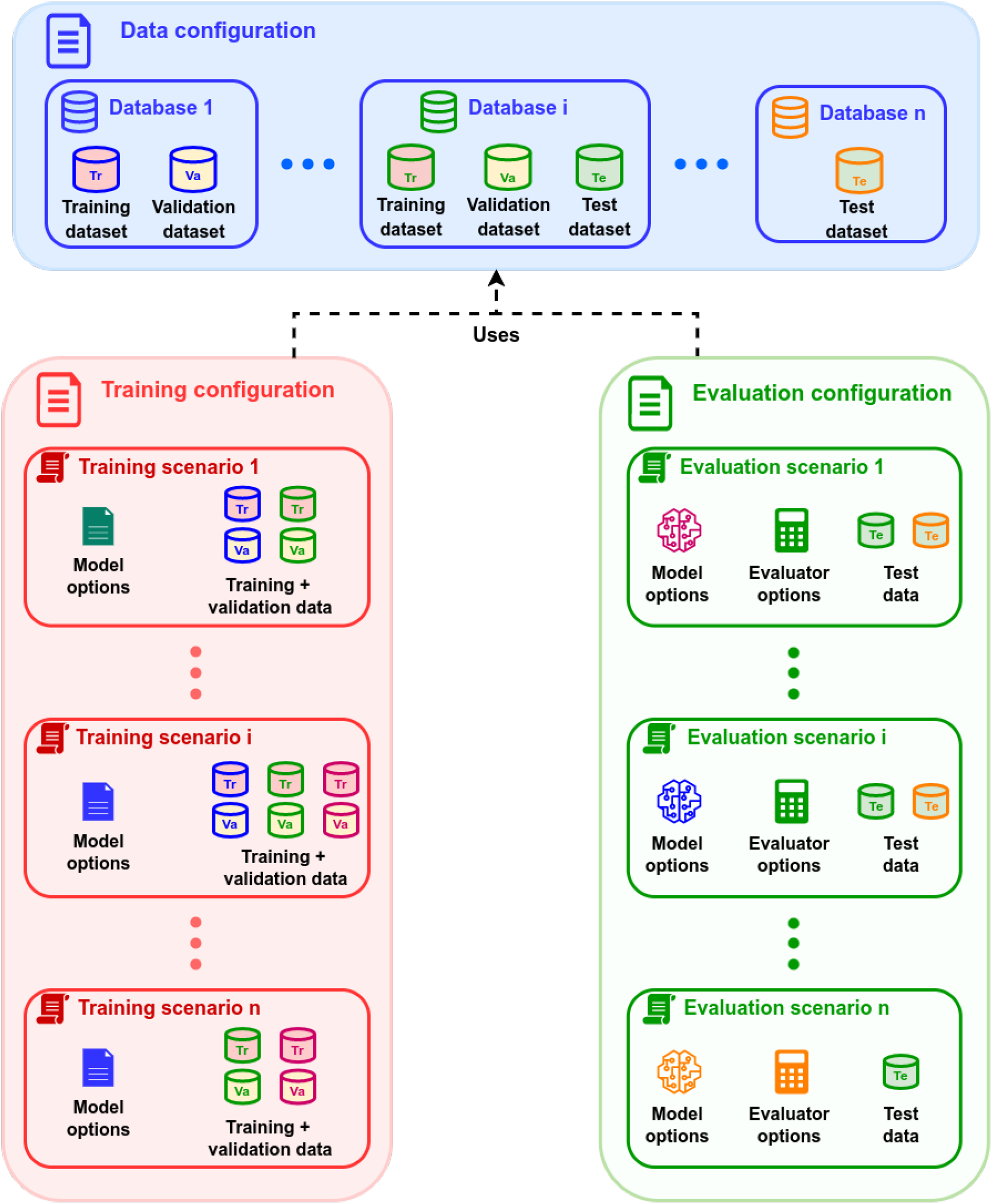
Organisation and contents of the three configuration files required by Mouffet. Datasets used in training and evaluation files should be defined in the data configuration file.

#### Data configuration file

This file contains all information related to data, including paths to where the data is located and how it is organized. Mouffet structures data on two levels: databases and datasets. A database is a collection of data sharing a common theme. It could be, for instance, data coming from the same source or collected in the same location. A dataset is a subdivision of a database created for a specific model creation task. We identify three types of datasets (for more details, see our previous paper (Christin et al., 2021)):

- Training dataset: used for training the model.
- Validation dataset: used for validating the model when the training is iterative. Usually, a subset of the training dataset
- Test dataset: used for evaluating the model. Should reflect the intended use case of the model.

#### Training configuration file

This file contains all options related to the creation of a model, including the paths where models will be saved, the databases, and the options that will be used to train a model. This file requires a link to a data configuration file.

#### Evaluation configuration file

This file contains all options related to the evaluation of a model, including the paths where the predictions and results of the evaluation will be saved, and the models, databases, and evaluators and their options to use for the evaluation. It also contains a reference to the data configuration file.

### 2.2 Scenarios

The main feature of Mouffet is the ability to define multiple scenarios in a single configuration file. A scenario is defined as a set of options that defines a task for a single model. For instance, a training scenario encompasses all the options required to successfully train a model.

Being able to define multiple scenarios allows the user to specify in a single file different values of a variable of interest, e.g., training parameters or detection thresholds. Behind the scenes, Mouffet looks for the key option “scenarios” and creates multiple configuration files, one for each of the scenarios (for a detailed example, see Fig. 3 below). Inside a ‘scenarios’ option, Mouffet search for lists of other options. Then for each list found, Mouffet creates all possible permutations of the values inside the list.

As the “scenarios” option can be present anywhere, virtually all the variables defined in the configuration files can be changed within a scenario; the only exceptions being a small set of variables required by Mouffet, mainly paths to other configuration files or where to save data. To keep track of what happens and to improve reproducibility, all the options used for a single scenario are saved for each task.

### 2.3 Runs

By default, training and evaluation tasks are processed separately. During evaluation, it is therefore required to manually define which model to evaluate. Since this can become complicated when multiple models are involved, Mouffet introduces the notion of runs.

Runs are simply sets of scenarios where one or several models will be both trained and evaluated. This means that all models defined in the training configuration file will be automatically evaluated using the options defined in the evaluation configuration file. Runs require a complete set of configuration files to work (data, training, and evaluation). For added convenience, training can be skipped if a model already exists, meaning that a run can be interrupted or repeated without having to train the models each time.

### 2.4 Python implementation

The Mouffet package provides all the logic for handling the configuration files and running the different scenarios. However, since models and databases can differ between users, Mouffet does not provide any logic on how to handle the data or how to perform the training or evaluation tasks. Instead, Mouffet defines Python objects that users must implement according to their needs using the Python inheritance system. Each component has access to the relevant configuration files, allowing users to define their own options. Here we describe the most important classes to implement for the framework to run properly. The general workflow of the package can be found in Fig. 2.

**Figure 2:**
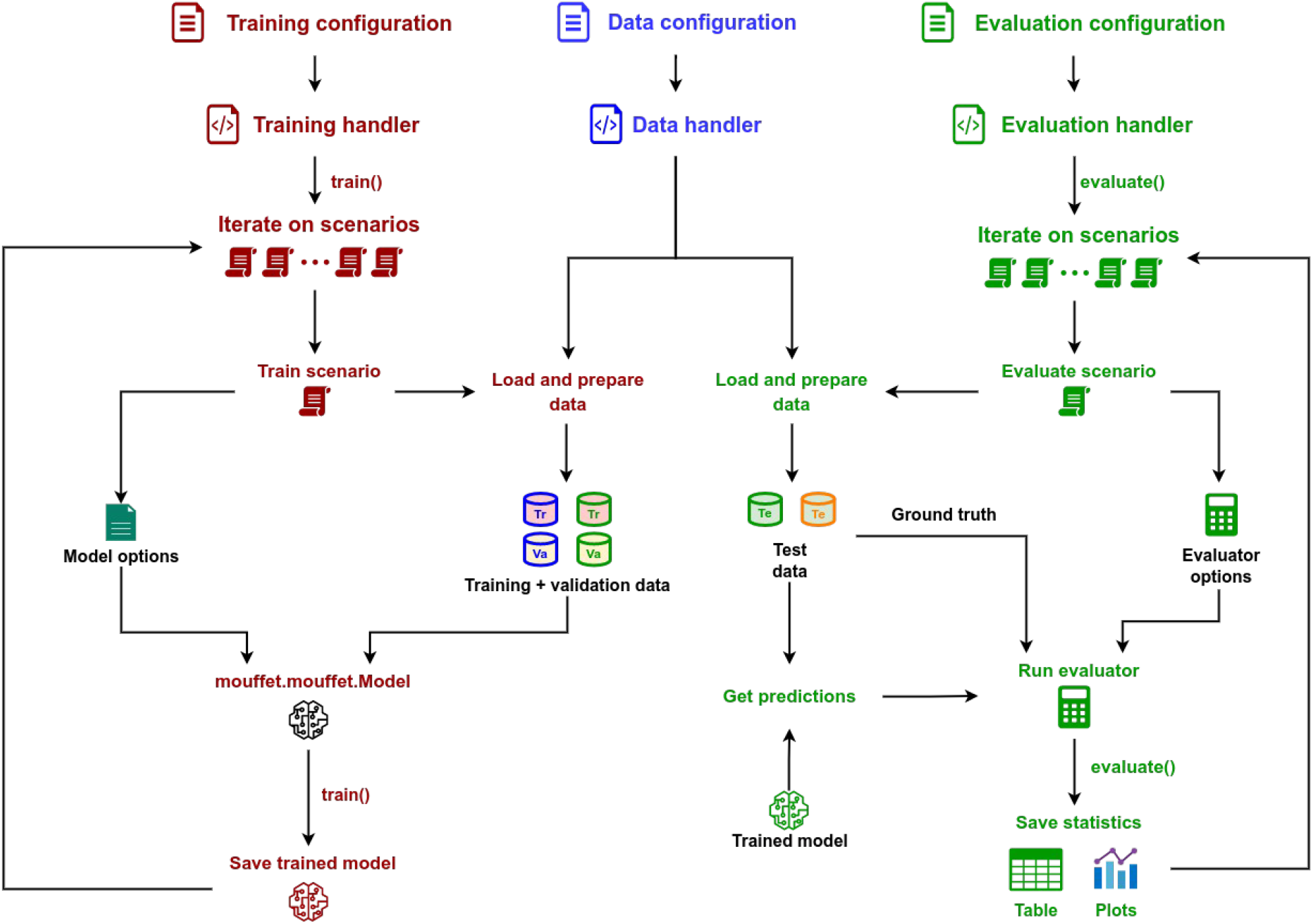
General training and evaluation workflow of Mouffet.

#### 2.4.1 Dependencies

Mouffet is written in Python v3.8 and requires the Python Standard Library (https://www.python.org). The package is lightweight and initially requires only three additional dependencies: “pandas” for dataframe manipulation (Reback et al., 2022), “pyyaml” for reading YAML configuration files (https://pypi.org/project/PyYAML/) and “feather-format” (https://pypi.org/project/feather-format/) to save dataframes in the feather format, a fast and lightweight saving format designed to be easily used with the R programming language.

#### 2.4.2 Data management

##### a) Data representation

In Python, data is represented by two classes: mouffet.data.Database and mouffet.data.Database.

The ‘Database’ class stores information about a database, such as the paths to where data is located and provides functions to manage datasets such as checking if a dataset exists or splitting the data into the appropriate datasets.

The ‘Dataset’ class represents a dataset. It describes how the data is organized in Python, i.e., where to find, for instance, raw data, annotations, or metadata. This class also provides convenience functions for loading and saving datasets.

##### b) Data manipulation

Data is handled with an implementation of the mouffet.data.DataHandler class. This class mainly helps manage databases and datasets. One of its most important tasks is to prepare datasets for training or evaluation, i.e., modify the raw data to fit the models. This could include data augmentation or normalization, e.g., by resizing images to make sure they all have the same size. Any modification to a dataset should be done in the *prepare_dataset()* method.

#### 2.4.3 Models

To define a model, users must provide an implementation of the mouffet.models.Model class. The model classes are abstract and will not work without a custom implementation of some key methods that vary depending on the type of model. Essentially, the user will have to define how to create, load/save, and use the model. As Mouffet was initially designed with deep learning in mind, Mouffet also provides the mouffet.model.DLModel class for this type of model.

#### 2.4.4 Training

Training in Mouffet is handled by an instance of the class mouffet.training.TrainingHandler. This class handles the logic related to the training options and expands all training scenarios. For each scenario, it loads the appropriate model, and then calls the train() method of the model with the appropriate data. During training, only training and validation datasets will be used. Each model is then saved, as well as the options that led to its creation.

#### 2.4.5 Evaluation

##### a) Evaluation handler

The evaluation handler is defined by the mouffet.evaluation.EvaluationHandler class and provides functions to load and expand evaluation scenarios from the evaluation configuration file. For each scenario, it loads the required data, the model, and calls the model to predict the data. It then calls an evaluator to calculate the desired metrics on the predictions. Note that a single scenario uses only one model, one evaluator, and one test dataset. By default, all predictions are saved and reused if a scenario is run again. Once all scenarios have been run, results are compiled and saved with each set of options. The evaluation handler also offers the possibility to generate plots to describe all the results that will be saved in a pdf file.

##### b) Evaluator

The mouffet.evaluation.Evaluator class takes a set of predictions made on a test dataset, compares them with the expected results, and then calculates evaluation metrics. Each evaluator should represent a different way of evaluating the predictions. It is possible to generate plots for the evaluator. All plots for scenarios with a similar evaluator will then be compiled in a single multi-page pdf file.

## 3. Example: classifying flower images with Tensorflow

To show the capabilities of Mouffet, we adapted a data augmentation tutorial from the Tensorflow website (https://www.tensorflow.org/tutorials/images/data_augmentation). That tutorial aims to develop a simple deep learning model to classify images of five types of flowers using data augmentation techniques such as flipping and rotating the images. Data augmentation is a common practice in deep learning to increase the number of available examples, reduce overfitting, and increase training accuracy (Shorten & Khoshgoftaar, 2019). After training, for each class of flower, the model gives a probability that an image belongs to this class. Model performance is then assessed by selecting the class with the highest probability and comparing it with the ground truth.

In this example, we adapted that model using Mouffet’s workflow to train five different models using each different parameters of data augmentation. We defined a custom evaluator that allows the user to choose a probability threshold for an image to be deemed well classified. This means that, if the classification probability of the best class is under the selected threshold, the image will be assigned the label “Unsure”. Mouffet’s plotting capabilities are used to compare the performance of each model with different tolerance thresholds and to generate confusion matrices for each evaluation scenario.

Please note that this example aims only to showcase Mouffet features and potential; it is not a deep learning tutorial, nor a fully-fledged research example. Since the concepts behind Mouffet extend beyond deep learning, no prior knowledge of working with Tensorflow or even deep learning, is required. We, however, assume that users have a basic knowledge of Python. All steps describing how to run the example can be found at https://github.com/Vin985/mouffet/blob/main/EXAMPLE.md

### 3.1 Dependencies

For the following example, the following additional libraries will be required: “tensorflow” (Martín Abadi et al., 2015) “tensorflow-datasets”, “scikit-learn” (Pedregosa et al., 2011) (to easily calculate evaluation metrics), and “plotnine” (https://plotnine.readthedocs.io/en/stable/) (a plotting library implementing the grammar of graphics used in the ggplot2 R package).

### 3.2 Data management

The data used in this example is taken from the”tf_flowers” database that can be downloaded and accessed via the “tensorflow-datasets” package. This database contains 3,670 images of flowers separated into 5 classes (dandelion, daisy, tulips, sunflowers, and roses).

The data configuration file for this example is simple. We defined a database named “tf_flowers” in the “databases” options. Since the data will be downloaded from the internet, there is no need to define paths here. For our demo example, we only defined how the datasets are split and chose to put aside 80% of the dataset for training, 10% for validation, and 10% for testing.

Python-wise, in the file data.py, we defined a ‘FlowersDataset’ class that loads the dataset using the tensorflow interface. We also created a ‘FlowersDatabase’ class that bypasses default functions from Mouffet as the package assumes and checks whether data is present on your hard drive.

The most important feature is the prepare_data() function found in the FlowersDataHandler class from the evaluation_handler.py file. This is where data augmentation for the dataset occurs, i.e., where images are rotated and flipped the images. Values that define how much to rotate the image and how to flip it are taken from the configuration file. For more details about the implementation, please refer to the user guide found online at https://mouffet.readthedocs.io/en/latest/tutorial.html.

### 3.3 Model

We created a FlowersClassifier class that essentially copies the tutorial o define the model in the create_model() function. We also defined the train() function and added logic for early stopping, a practice that stops the training automatically when no more progress is made. Once again, this is customizable from the training configuration file. Finally, we specified how to load/save and use the model.

### 3.4 Training

#### 3.4.1 Training configuration

Here, besides the paths to the data configuration file and where to save the models, we defined global options for the models such as the class and database to use.

Then we set five training scenarios, described in the “scenarios” option (Fig. 3).

**Figure 3:**
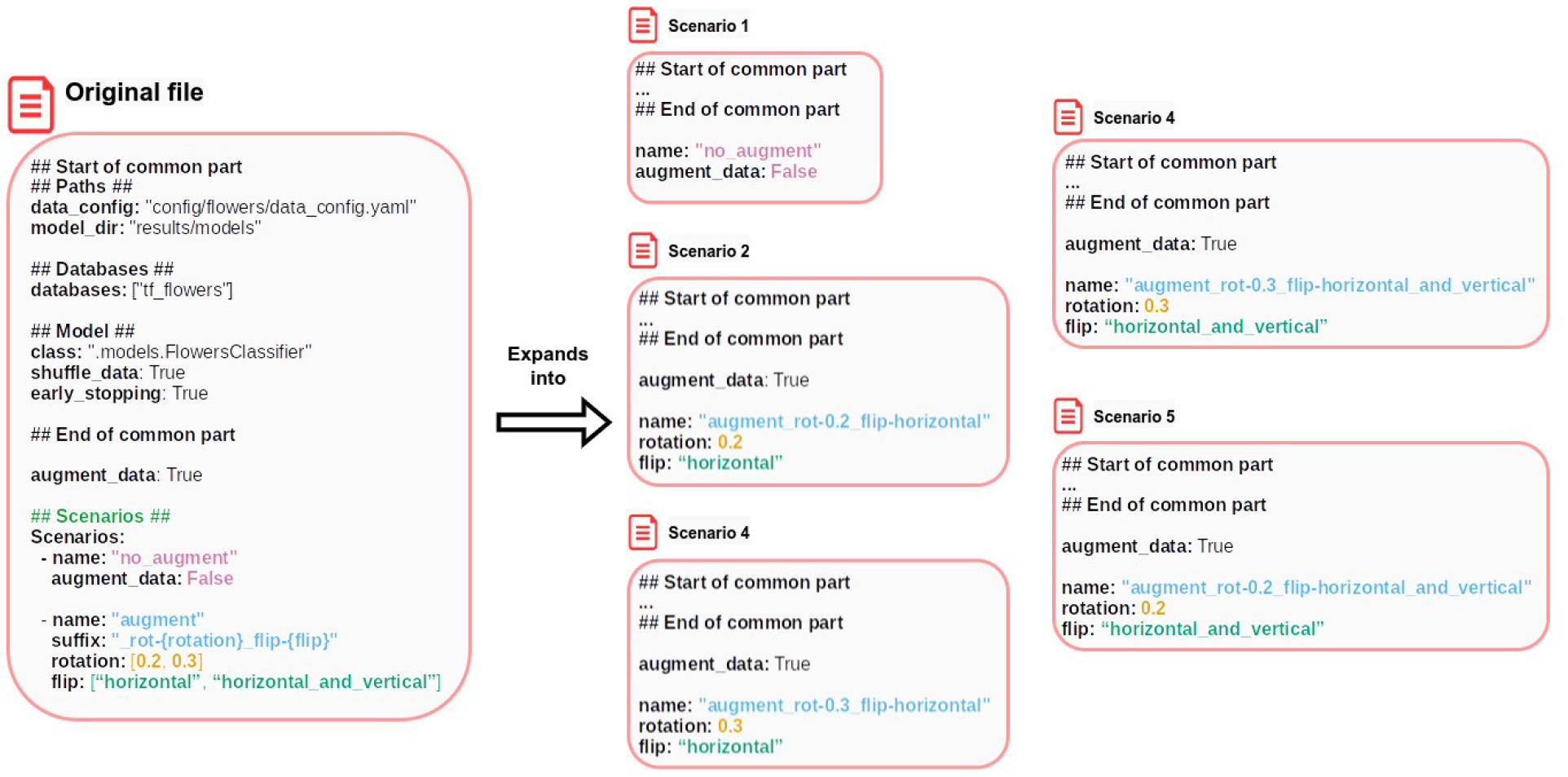
Example of scenario expansion of a configuration file in Mouffet. On the left, the original file with scenarios defined in it. Upon expansion, Mouffet creates five configuration files with only some options changing. All the rest remains the same between files. When a list is present, such as in the ‘rotation’ and ‘flip’ options, all possible permutations are generated. Note the presence of the ‘suffix’ option in the second entry of the scenarios. This option allows for a dynamic model name generation. Every string between brackets {} will be replaced by the value of the option of the same name. The suffix is then appended to the ‘option’.

**Figure 4:**
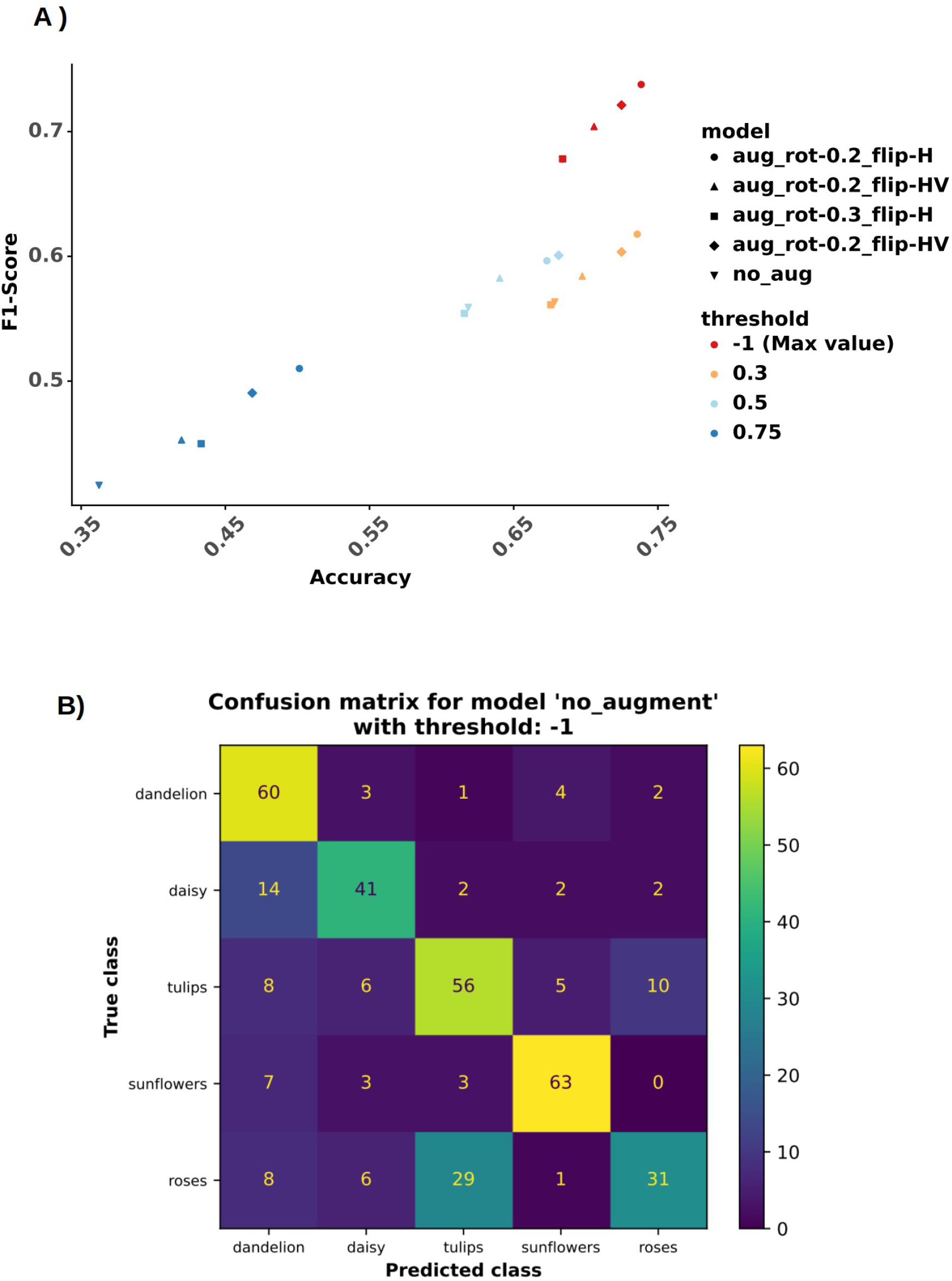
Plots generated by Mouffet during the evaluation of the trained flowers classifiers. A) F1_score as a function of the accuracy of the trained models with different thresholds selected during the evaluation. B) Example of a confusion matrix generated by the evaluator. Confusion matrixes are generated for each evaluation.

- First, we defined a scenario with no data augmentation where the “augment_data” option is overriden (Fig. 3 – scenario 1).
- Then we defined augmentation scenarios where we changed parameters related to the rotation and flip. We set two lists of values: one for the ‘rotation’ parameter to take values 0.2 and 0.3 (the maximum amplitude on which to rotate the images) and one for the ‘flip’ option to take values ‘horizontal’ and ‘horizontal_and_vertical’. During training, Mouffet iterates over these lists and creates all the permutations possible for these values. This results in the same as defining 4 models with different values for rotation and flip. To keep track of these values, we added a suffix to our model’s name. Each variable name between brackets will be replaced by the actual value of the variable during the training (Fig. 3 – scenarios 2-5).

#### 3.4.2 Training the models

Training occurs in the training.py file. Since the example here is quite simple, we did not need to implement a new training handler and used the one provided with Mouffet instead.

We just created the TrainingHandler and called the train() method. We then obtained 5 models that can be found in the folder specified in the ‘model_dir’ option.

### 3.5 Evaluation

#### 3.5.1 Evaluation configuration

Once more, we defined paths where to find models, and databases and where to save the results. Three options are essential here:

- ‘databases’: defines which databases will be used.
- ‘models’: defines which models will be evaluated. Here we simply copy the training configuration file.
- ‘evaluators’: defines the evaluators to be used. Each evaluator needs a unique name that will be defined during registration (See below for more information). Here we set 4 different values for the ‘threshold’ option inside a ‘scenarios’ keyword: -1, 0.3, 0.5 and 0.75

With the ‘models_options’ and ‘evaluators_options’, we defined options that will be applied to all models and evaluators respectively. Here we did not want to augment the data and told the evaluators to plot confusion matrixes. We also defined global plots that will be generated from all the results, not just at the evaluator level.

#### 3.5.2 Evaluator

To implement the behaviour described above, we created a subclass of mouffet.evaluator.Evaluator called CustomEvaluator in the file evaluators.py. In its *evaluate()* method we added a ‘threshold’ option taken from the configuration file that behaves like this:

- If ‘threshold’ == -1, the evaluator behaves like the original tutorial, i.e., the selected class is the one with the highest given probability.
- If 0 <= ‘threshold’ <= 1, and the highest probability is above the threshold, its associated class is returned. Otherwise, the image is attributed to the class “Unsure”.
- Otherwise, an error is raised.

We used the method ‘classification_report()’ from the package scikit-learn to calculate metrics such as precision, recall and f1-score for each class and globally. We also added the plot_confusion_matrix(), which plots confusion matrices using the scikit-learn package.

#### 3.5.3 Evaluation handler

In the file evaluation_handler.py, we created a custom evaluation handler called FlowersEvaluationHandler inheriting mouffet.evaluation.EvaluationHandler. In this class, we told Mouffet how to generate predictions from the test dataset in the predict_database() method. We also implemented the function plot_accuracy_f1() to plot the f1_score as a function of the accuracy for all the results.

#### 3.5.4 Performing the evaluation

Evaluation is done in the evaluation.py file. The concept is the same as for training, we created the FlowersEvaluationHandler and just called the evaluate() function. The main difference is that we needed to register our evaluator by giving it a unique name to be able to use it. That name is the one put in the configuration file in the ‘evaluators’ option. Once the evaluation was done, we obtained the following results:

- In the results/predictions folder, the predictions for each model on each database. were saved in the feather format. A predictions_stats.csv files that compiled information about how the predictions were generated (time spent, options used, etc.) was also created.
- In the results/evaluation folder, we got the evaluation results. Here this consists in: a csv file that contains all results - including the options used to generate them - in a table format; a pdf with all confusion matrixes (figure 2-B) plots and a pdf with the accuracy/f1 score plots generated from all results (figure 2-A). Note that all files are time-stamped and sorted by date to keep track of when the evaluation was performed.

### 3.6 Launching a run

To avoid having to launch separately the training and evaluation, we can define a run that will do both for us. Runs are named, and all the configuration files should be put inside a folder with the run name. Here we put them in the ‘flowers’ folder. Then inside the file run.py, we created a RunHandler that takes all handlers as arguments. We then called the run.py file by passing the name of our folder as a command line argument. We obtained the same results as if we launched the processes manually, except that results are now saved in the results/runs/flowers folder.

## 4. Discussion and outline

Here we introduced Mouffet, a framework that aims to make model creation easier, more robust, and more reproducible. By defining multiple training and evaluation scenarios in a single configuration file, users can create experimental protocols to evaluate the impact of a set of variables on model performance. Besides, by saving in a structured way all results, along with the options used to create them, it becomes easier to repeat these experiments.

However, while Mouffet offers tools to assist model creation, some good practices remain up to the users. To improve reproducibility, while multiple scenarios can be defined in a single file or for a given run, we recommend creating new sets of configuration files for each iteration of the model development instead of just updating the same ones. For instance, if we want to study the influence of image resolution and data augmentation on the performance of an image classifier, we could create a set of configuration files that modify the resolution and another set centered on data augmentation. This would help keep track of the progress made during the creation process, which in turn makes it easier to reproduce and share.

Yet, while this provides a way to track the evolution of options, this does not apply to the underlying implementations of the models. Creating new models each time could be an option also but could soon become cumbersome as models become obsolete. Instead, we strongly suggest using some sort of version control software, such as git (Chacon & Straub, 2014) to keep track of changes in the code itself. Ultimately, it remains up to the users to make the most of Mouffet’s capabilities to ensure that their model creation process can be as reproducible as possible.

Since Mouffet is independent of the underlying implementation of the models, it should work with most existing model frameworks. And while it was developed with deep learning in mind, its concepts also apply beyond this domain. Therefore, it should be possible to create any kind of model, especially other machine learning approaches. This makes Mouffet an ideal tool to compare different modelling approaches.

Thanks to its versatility and ease of use, especially for non-computer scientists, we believe Mouffet could become a choice tool for ecologists looking to develop models in a robust and reproducible way.

## Acknowledgements

The study was funded by the Canada Research Chair in Polar and Boreal Ecology and the New Brunswick Innovation Fund. The authors declare no conflict of interest.

## Author contributions

S.C. designed and implemented Mouffet with input from N.L. S.C. and N.L wrote the first draft of the manuscript and N.L. provided comments.

## Data availability

All code is freely available at https://github.com/Vin985/mouffet

